# MetQy – an R package to query metabolic functions of genes and genomes

**DOI:** 10.1101/215525

**Authors:** Andrea Martinez–Vernon, Frederick Farrell, Orkun S. Soyer

**Affiliations:** Synthetic Biology Centre for Doctoral Training, University of Warwick, Coventry, CV4 7AL, UK; School of Life Sciences, University of Warwick, Coventry, CV4 7AL, UK; Warwick Integrative Synthetic Biology (WISB) Centre, Life Sciences Building, University of Warwick, Coventry CV4 7AL, UK.

## Abstract

**Summary:** With the rapid accumulation of sequencing data from genomic and metagenomic studies, there is an acute need for better tools that facilitate their analyses against biological functions. To this end, we developed MetQy, an open–source **R** package designed for query–based analysis of functional units in [meta]genomes and/or sets of genes using the The Kyoto Encyclopedia of Genes and Genomes (KEGG) database. Furthermore, MetQy contains visualization and analysis tools and facilitates KEGG’s flat file manipulation. Thus, MetQy enables better understanding of metabolic capabilities of known genomes or user–specified [meta]genomes by using the available information and can help guide studies in microbial ecology, metabolic engineering and synthetic biology.

**Availability and Implementation:** The MetQy R package is freely available and can be downloaded from our group’s website (http://osslab.lifesci.warwick.ac.uk) or GitHub (https://github.com/OSS-Lab/MetQy).

## 1 Introduction

The advent of molecular biology has made the characterization and analysis of genomic sequences a key part of all areas of life sciences research. In the case of single–cell organisms, identification of specific functions within the genome directly influences our ability to assess their fitness in a given environment and their potential roles in biotechnology. Particularly, we should theoretically be able to translate genomic data into physiological predictions. Genomic databases are a prerequisite for making such predictions, but their full use also requires computational tools that allow easy access and systematic analyses of the data.

The Kyoto Encyclopedia of Genes and Genomes (KEGG) is one of the oldest and most comprehensive databases. Its primary aim has been the digitizing of current knowledge on genes and molecules and their interactions (Kanehisa, 1997; Kanehisa and Goto, 2000) and it includes 16 databases and 3 sequence data collections (Kanehisa *et al.*, 2017). While these data can be analyzed via different tools on the KEGG website, the existing web interface allows only specific retrieval of information and analyses. Furthermore, although the whole of the data can be downloaded via (paid) FTP access, the systematic analysis of these data in a user–defined manner remains difficult and developing computational analysis tools for this purpose remains a niche expertise that is still not available in many research labs.

There are several specific tools that make use of certain aspects of the KEGG data more available to a wider user-base. Examples include PICRUSt (Langille *et al.*, 2013), BlastKOALA and GhostKOALA (Kanehisa *et al.*, 2016), all of which focus on metagenomics data analysis. However, to our knowledge there are no tools that facilitate the analyses and information retrieval from KEGG with regards to studying the relationship between genomic data and physiological function. Therefore, we have developed MetQy, an open–source, easy–to–use and readily expandable **R** package for such analyses. In particular, the tool allows analysis involving so-called KEGG modules, collections of genes associated with well–defined metabolic and other functions. MetQy uses the **R**–platform because it is commonly used among biologists, it is featured in undergraduate education, and it contains extensive statistical packages which are useful in subsequent data analyses.

MetQy was developed to readily interface between the KEGG orthology, module and genome databases and perform automated cross–analyses on them. It consists of a set of functions that allow querying genes, enzymes and functional modules across genomes and vice versa, thereby enabling better understanding of genotype–phenotype mapping in single–celled organisms and providing guidance for cellular engineering in synthetic biology. The MetQy package contains extensive documentation and usage examples for each function.

## 2 Software Features

MetQy contains three main groups of software functions: data query, parsing, and analysis and visualization. These are briefly described below. For more detailed information and usage examples, please see the package documentation and GitHub wiki.

### 2.1 Metabolic query functions

The *query* family of functions allows the user to query the KEGG data structures in a systematic (and automated) way. Users without FTP access can analyze the KEGG genome, module and ortholog databases indirectly by using this family of functions on built–in formatted KEGG data (downloaded in July 2017) which is not directly accessible by the user. Additionally, these functions feature optional arguments that allow users to provide up–to–date data (by using the *parsing* functions on KEGG FTP data) or their own data structures, such as custom–made KEGG–style modules or metagenomes. Additional query functions can be readily developed by the users, allowing expansion of MetQy. MetQy features five query functions for key functional analyses.

*query_genomes_to_modules* calculates the module completeness fraction (*mcf*) given a set of genes or genomes. It returns a matrix showing the *mcf* for each module. The *mcf* calculation is based on block– based, logical KEGG module definition (see https://github.com/OSS-Lab/MetQy/wiki/KEGG-databases-description). The function input is the modules to be queried (default is all KEGG modules) and the set of of genes to be considered provided either as a set of KEGG ortholog or Enzyme Commission (EC) numbers or as genome identifier(s), with the latter case resulting in automatic retrieval of all genes for the genome(s).

While the implementation of “*query genomes to modules*” function is similar to KEGG mapper (a web interface tool that performs a similar task (http://www.genome.jp/kegg/mapper.html; Kanehisa *et al.* (2017))), there are several key features that are different. First, the KEGG Mapper’s web interface does not allow for module–specific evaluation nor for automation of the analysis. Our implementation allows for specific KEGG modules to be evaluated, given their ID, name and/or class. It also provides the capacity to determine the *mcf* of a module, rather than only identifying modules that are complete or that have one block missing. Finally, as EC numbers are widely used in systems biology, we used the KEGG orthology to translate the K number–based module definitions to EC number–based module definitions. This allows for module evaluation based on both K and EC numbers.

*query_module_to_genomes* determines the KEGG genome(s) that have user–specified module(s) that are complete above a *mcf* threshold (defaults to 1, i.e. complete). The function output is a matrix showing *mcf* values organized across module IDs (columns) and genome identifiers (rows).

*query_gene_to_modules* determines those KEGG modules that feature specific user–specified gene(s).

*query_genes_to_genomes* determines which KEGG genomes contain user-specified gene(s).

*query_missingGenes_from_module* determines the missing gene(s) (K or EC numbers) that would be required to have a complete KEGG module within a genome (or gene set).

### 2.2 Parsing KEGG databases

The *parsing* functions allow users with current KEGG FTP access to increase the usability of the KEGG databases and to provide up–to–date data structures to the *query* family of functions. MetQy features two generic parsing functions that deal with the two main KEGG file types: files without extension (*parseKEGG_file*) and ‘.list’ files (*parseKEGG_file.list*). MetQy also contains file–specific functions that use these generic functions.

*parseKEGG_file.list* formats KEGG files containing a mapping between two KEGG database entries into binary matrices. For example, the mapping between K numbers and EC numbers is contained in the ‘ko enzyme.list’ file and shows which K numbers correspond to which EC numbers.

*parseKEGG_file* formats a KEGG database file into an **R** data frame by automatically detecting fields of the KEGG data and transforms these into variables.

### 2.3 Analysis and visualization

The analysis and visualization family of functions is designed to facilitate the analysis of the output of the *query* functions. They have the prefixes *analysis*_ and *plot*_, respectively. The former includes variance and principal component analyses, while the latter includes custom–made heatmaps and various scatter plots.

## 3 Uses and applications

MetQy facilitates the general usability of the KEGG database and allows users to gain qualitative information about the functional capacity of a given organism or gene set. Anticipated uses of the tool include synthetic biology, where it can facilitate the design and guiding of metabolic engineering studies by identifying missing genes needed for an organism to have a complete KEGG module, and identifying KEGG genomes with desired metabolic capabilities. For systems biology applications, it allows identification of key physiological features of organisms and development of stoichiometric metabolic models by analyzing module completeness in specific genomes and identifying transporter modules and carbon utilization routes in genomes. Finally, in microbial ecology, MetQy can allow species-function mappings in metagenomes and insights into functional capabilities of ecological groups by analyzing the metabolic capacity of novel genomes from metagenomic studies. Organisms can be put into different functional groups, and the functional profiles of different environments compared.

## Acknowledgements

We acknowledge Sean Aller for helpful comments and David Selby for sharing his expertise in developing **R** packages.

